# What difference do retractions make? An estimate of the epistemic impact of retractions on recent meta-analyses

**DOI:** 10.1101/734137

**Authors:** Daniele Fanelli, David Moher

**Affiliations:** Department of Methodology, London School of Economics and Political Science, London, UK; Centre for Journalology, Clinical Epidemiology Program, Ottawa Hospital Research Institute; School of Epidemiology and Public Health, University of Ottawa, Ottawa, Canada

## Abstract

Every year, several hundred publications are retracted due to fabrication and falsification of data or plagiarism and other breeches of research integrity and ethics. Despite considerable research on this phenomenon, the extent to which a retraction requires revising previous scientific estimates and beliefs – which we define as the epistemic impact - is unknown. We collected a representative sample of recently retracted studies that had been included in recent meta-analyses, and compared the summary effect size of these meta analyses with and without the refracted studies. On average, the retractions had occurred about six years prior to the publication of the corresponding meta-analyses.

Our results suggest that retractions have varying impacts depending on their causes. In particular, removing from an analysis a study retracted because of issues with data, methods or results, led to a statistically significant reduction of the estimated effect size. Assuming that the results of these retracted studies are completely false, then the meta-analyses that had included them had overestimated the summary effect sizes by, averaging across effect size metrics, 30% (median, 13%). However, retractions due to plagiarism or other issues not related to data, methods or results had no impact on the conclusions of meta-analyses.

Since retractions due to plagiarism or other non-data related issues typically constitute over 75% of total retractions, our results suggest that the epistemic impact of most retractions is likely to be null. However, our results also suggest that retractions due to issues with data, methods or results should be accompanied by a revision of relevant meta-analyses, and by extension a downwards revision of prior scientific beliefs.

---

Retractions are a phenomenon of growing importance in science, yet it remains unclear if and to what extent they constitute an obstacle to scientific knowledge, rather than a positive manifestation of scientific self-correction.

The number of retracted publications has grown from practically zero three decades ago to at least 1,161 items labelled as “retraction” in 2018 in the Web of Science database alone. However, contrary to common concerns that misconduct is rising in science, multiple lines of evidence suggest that the growth in retractions results mainly or entirely from the expansion and strengthening of policies and practices to correct the literature [1]. If retractions manifest scientific self-correction, then their recent rise is to be celebrated as a positive development that should be further encouraged [2,3].

Multiple independent studies have documented how the authors of a retracted study suffer a significant cost in terms of citations, productivity and funding [4–6]. However, these costs are only observed for authors of studies retracted due to misconduct. Authors of articles retracted for honest error appeared to suffer no negative consequence [4] and collaborators of authors of retracted articles suffer a loss of citations mainly when the retraction was due to misconduct [7–9]. If citations and career costs affect exclusively individuals who commit scientific misconduct, then they are not “costs” at all, but fair sanctions administered collectively by the scientific community, which act as deterrents and should be supported.

A clearer case may be made that retractions entail a waste of resources. However, the financial costs are surprisingly contained. A recent study, in particular, estimated the financial impact of studies that were funded by the US National Institutes of Health and were retracted due to findings of misconduct. It concluded that, even using approximations that over-estimated them, the costs amount to between 0.01% and 0.05% of the NIH budget, a figure that was deemed low in the authors’ own assessment [10]. Furthermore, retracted studies constitute wasted resources only to the extent that their results are technically invalid, which is not obvious to be the case even when retractions are due to fabricated data, since the claims made by a fabricated study could be scientifically correct [11].

Several studies have documented the fact that retracted articles keep being cited after their retraction, although typically at much lower rates [5,6]. However, part of these post-retraction citations are likely to be negative or neutral (for example, they cite the retracted article as an example of retraction). According to a small analysis of 15 cases, the majority of citations to retracted articles are positive, as if the retraction had not occurred [12]. If confirmed on a larger scale, this could represent a disturbing phenomenon, which may result, in part, from flaws and limitations in the electronic indexing systems of literature databases, which render citing authors unaware of the retracted status of an article [13] and, in part, from authors who wilfully ignore the retracted status of an article. However, whereas citing a retracted article is unquestionably problematic from an ethical point of view, because it undermines the sanctioning role that a retraction ought to have on its authors, from a scientific point of view the costs are less immediately obvious. The argument is similar to that made above for financial costs: citing a retracted article is harmful to scientific knowledge only to the extent that the retracted article reports incorrect or distorted knowledge.

The extent to which scientific knowledge is actually distorted is what we will call the “epistemic cost” of a retraction. The magnitude of this cost has not yet been accurately estimated. Mata analysis offer a natural tool to measure this impact. If retracted studies entail no epistemic cost, removing them from a meta-analysis should have no impact on the meta-analytical results. Vice-versa, a difference will be observed if and to the extent that retracted studies reported distorted information. In a meta-assessment of bias in science, studies with a first author who had had other articles retracted were significantly more likely to over-estimate effect sizes [14]. Therefore, we hypothesize that meta-analyses that include retracted studies may over-estimate effect sizes.

A recent study examined the impact that a clinical trial that contained falsified data had on 22 meta-analyses that included it, and concluded that 10 (46%) meta-analyses had their results significantly changed[15]. This result, however, cannot be generalised due to various limitations. It examined a single instance of data fabrication, even though retractions can occur variety of reasons. Further, it included meta-analyses of the same all very similar research question. Furthermore, the fabrication of data was uncovered exclusively in a single Chinese clinical site and, once all Chinese data was excluded, the trial yielded statistically significant positive results [16]. Therefore, the average distorting effect that retractions exert on the scientific literature remains to be assessed.

This study estimates the epistemic cost of a representative sample of recent retractions, by measuring the difference that these make in the corresponding sample of meta-analyses that included them.

## MATERIALS AND METHODS

### Meta-analyses sampling strategy

We first compiled a list of all records that were tagged as “retracted” in the Web of Science database (WOS) as of 16 December 2016. These records are readily identified because their title in the WOS database is modified by the addition of the label “retracted article” or “retracted title”. The list was hand-inspected to exclude any false positives, leading to a total list of 3,834 records at the time (the WOS has recently updated its retraction tagging system and database, and now includes a large number of retractions).

We subsequently compiled a list of all records in the WOS that cited any of the retracted records, obtaining an initial list of 83,946 records. We then restricted the above list to records that included “meta-analysis” OR “meta analysis” OR “systematic review” in the title, abstract or keywords, which resulted in 1,433 titles. For each of these records, the full list of cited references was retrieved, and the one or more references that had been retracted was identified by matching the document object identifier (DOI).

Records for which the DOI did not identify any retracted cited reference were inspected by hand, and the corresponding retracted cited reference was identified by hand-searching the WOS.

We subsequently restricted the sample to an initial list of potentially usable meta-analyses published in 2016 and later expanded the sample to 2015. In both cases, we excluded records that, based on inspection of title and abstract, were identified as not being actual meta-analyses or systematic review. This left us with an initial list of potentially relevant records of N=109 for 2016 and N=120 for 2015.

### Inclusion/exclusion of potentially relevant meta-analyses

All but a few (N=9) of the Pdfs of the potentially relevant meta-analyses could be retrieved and manually inspected to determine exclusion based on the following criteria:

1. Is limited to a systematic review, and not a formal meta-analysis
2. Is not a standard meta-analysis, in that it does not produce a single weighted pooled summary of two or more primary studies. This excludes network meta-analysis, Genome-Wide-Association-Studies, meta-analyses of neuroimaging data, microarray data, genomic data, etc.
3. Does not contain a usable summary of primary data in the full-text or accessible appendix. In particular, we required it to give a funnel plot or table containing data on each primary study’s identity and reported effect size.
4. The retracted cited article is not one of the primary studies included in any of the weighted pool summaries presented.

The final number of included meta-analyses was N=31. Figure 1 summarizes the sample retrieval process in a flow chart.

**Figure 1.**
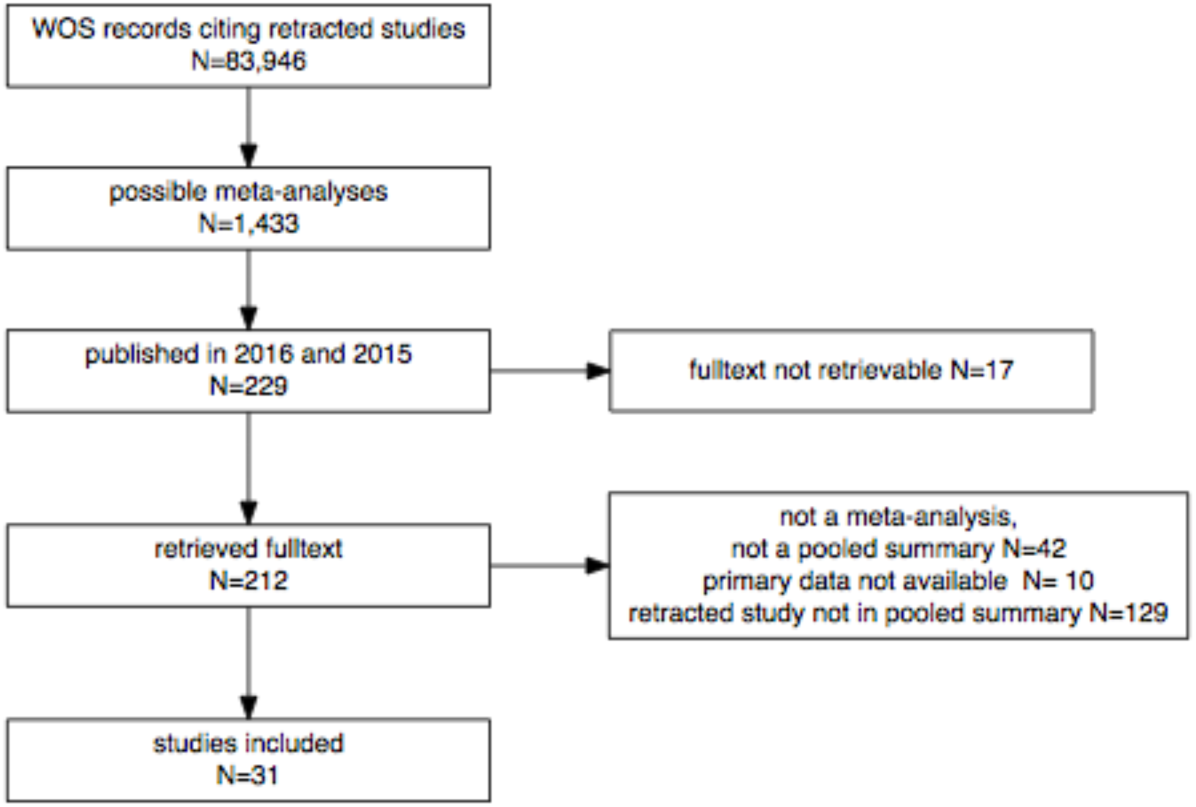
Flow chart of the sample retrieval process.

**Figure 2.**
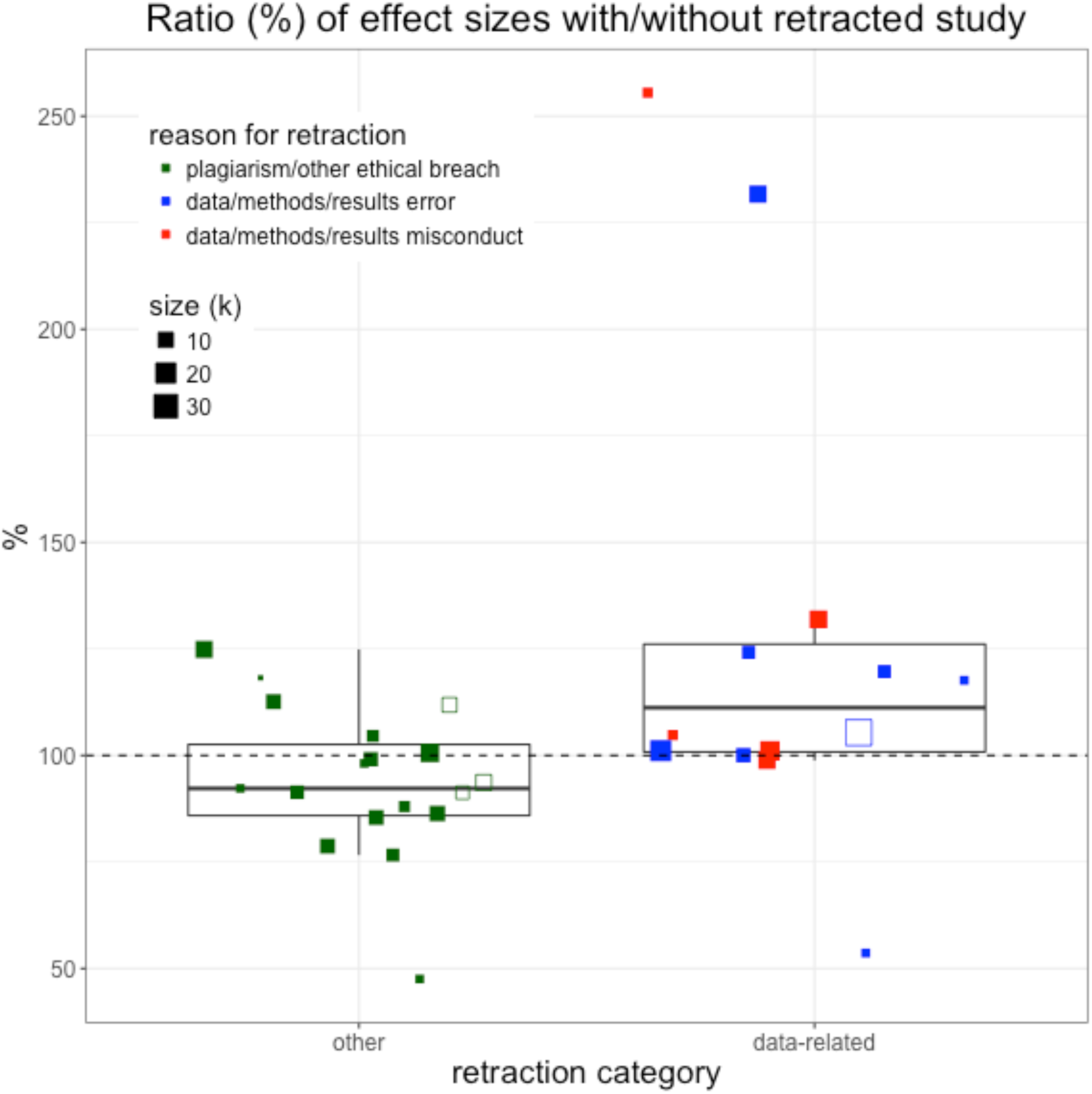
Relative size, in percentage, of meta-analytical pooled effect sizes calculated including and excluding the retracted study. Values of 100% (dashed line) indicate identical effect sizes, whereas values above 100% indicate that the meta-analysis with the retracted study yielded a larger effect size than without it. Empty squares indicate meta-analyses that included the same retracted study as another meta-analysis.

### Primary data extraction

From each included study, we identified the figure or table that reported the primary data for the meta-analysis. If the study contained more than one meta analysis, we selected the one that cited the retracted study, and if more than one such meta-analyses was present we selected the first one shown in the publication.

For each of the selected meta-analyses, we recorded the pooled summary estimate and for each of the primary studies within each meta-analysis, we recorded the reported effect sizes and confidence interval (or other measure of precision used, e.g. standard error, sample size)..

For each retracted primary study in our sample, we retrieved the text of the retraction note and we recorded the (one or more) reasons adduced in the note for the retraction. When in doubt about the reason, we compared the record with that in the Retraction Watch database (retractiondatabase.org).

### Calculating the impact of retractions

For each included meta-analysis, we calculated the summary effect size twice, i.e. of with or without the retracted primary study. Where possible, meta-analytical summary estimates were obtained using the primary raw data in the form used by the original meta-analysis (e.g. number of events and totals, or means and standard deviation and sample size, etc.). Alternatively, summary estimates were simply re-calculated based on the processed numerical data (e.g. standardized mean difference±95% confidence interval) reported in the figures (forest plots typically include data in numerical form) or table. Details of all calculations are given in the Supplementary Data File, and a comparison of original and re-calculated meta-analyses is given in the Results section.

To quantify the difference that removing the retracted study made to the meta-analysis, we adopted multiple strategies, because the included meta-analyses used different measures of effect size. In particular, with cultivated the following measures:

- Effect size ratio: ratio, expressed in percentage, of the effect size (ES) with vs. without the retracted study. For each meta-analysis *i*, this was calculated as:

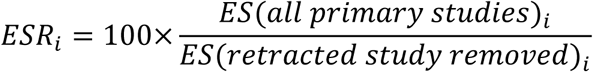

The ESR quantifies in very general terms how much larger or smaller the originally reported summary effect size is, compared to that corrected after a retraction. It therefore quantifies in the most general sense the impact that the retracted study had on the meta-analysis, regardless of the metric used by the meta-analysis. An ESR of 100 indicates no difference, and an ESR greater than100 indicates that including the retracted study led to a larger summary effect size in the meta-analysis.

To obtain a more metric-relevant estimate of the impact of a retraction, we limited our sample to meta-analyses with inter-convertible effect sizes (i.e. standardized mean difference, Hedges’ g, both assumed to be equivalent to a Cohen’s d, and Odds Ratio, Peto Odds Ratio and Risk Ratio, all three assumed to be equivalent to Odds Ratio). On the resulting sub-set of meta-analyses (N=13) we calculated the following metrics:

- ROR: ratio of odds ratios, which were inverted when necessary (IOR, see below), and calculated as:

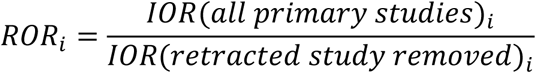

in which

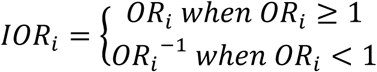

Is an appropriately “inverted” odds ratio. The inversion is necessary to align the (eventual) bias in an odds ratio-based meta-analysis.

- DSMD: difference between standardized mean differences (SMD). Indicating the latter as d, this is calculated as:

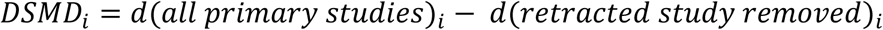
- DCC: difference in correlation coefficient, *r*, calculated as:

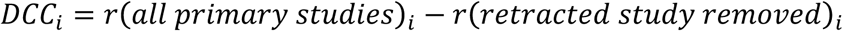

## RESULTS

### Descriptive results

In total, 31 meta-analyses could be included (14 published in 2015 and 17 from 2016). Table 1 reports the characteristics of these meta-analyses and the corresponding retracted studies.

**Table 1.**
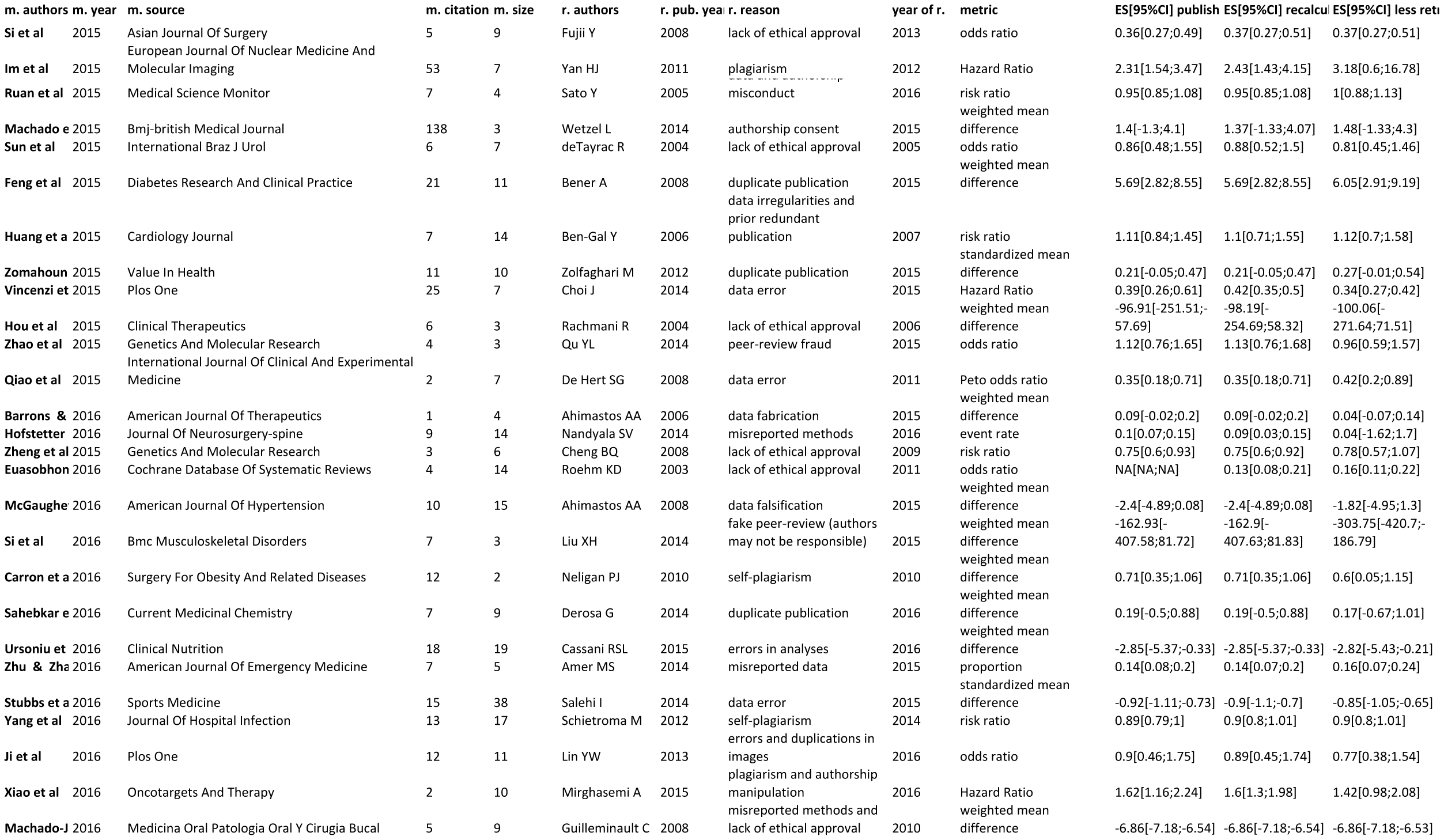
Characteristics of included meta-analyses, of their corresponding retracted studies, and effect sizes as reported in the original publication and as re-calculated by including or excluding the retracted primary study.

All but two of the included meta-analysis were published in clinical or biomedical journals, and the two exceptions, both published in the generalist open access journal PLoS ONE, were biomedical in scope nonetheless.

The included meta-analyses varied in their size (i.e. number of primary studies included, range: 2-38), in number of citations received (range: 1-138) and in the metrics they used. The most common metric used in the sample was weighted mean difference (n=12) followed by odds ratio (n=5) and risk ratio (n=4).

All of the retracted studies were unique to a single meta-analyses, with four exceptions (Bener 2008, Derosa 2014 and deTayrac 2004, Salehi 2014). The 27 unique retracted studies in our sample varied considerably in their year of publication (range: 1999-2015) and the year of their retraction (range: 2005-2016). On average, these studies had been retracted 3.4 years after their publication and 5.8 years prior to the publication of the meta-analysis that included them.

The reasons for retraction, as reported in retraction notices, covered a varied spectrum of possible ethical infractions, including lack of ethical approval, authorship or peer-review issues, various forms of plagiarism, errors in data or methods, and data fabrication and falsification. Although in some cases more than one reason for retraction was indicated, retractions could be reliably separated into two groups: those that were (partially or entirely) due to issues with data, methods or results (n=13) and those due to other causes (n=15).

### Recalculation of published effect sizes

In 30 of the 31 available meta-analyses, a pooled summary of the primary data set extracted had been calculated in the original publication (the exception was Euasobhon et al. 2016, in which the forest plot from which data was extracted was provided for a sensitivity analysis). The re-calculated effect sizes were generally in good agreement with values reported in the publications: in 25 cases, the re-calculated values were identical or within ±5% of the published value, and the average percentage deviation was 0.4%. The greatest discrepancy was found in Hofstetter et al. 2016, for which the re-calculated ES was 90.4% of the published value (i.e. event rate 0,09 versus 0,1). Any discrepancy between original and re-calculated value is likely to be due to details of the calculations (e.g. rounding of numbers, particular corrections) whose analysis was beyond the scope of the article. Because our objective was to assess the impact of retractions on pooled summaries, we used the re-calculated summaries as the estimate of comparison.

### Effects of retraction on meta-analytical summary effect sizes

The ratio effect sizes calculated with and without the retracted studies (henceforth, ESR,) varied substantially, ranging between 54% and 256%, and its distribution was non-randomly relative to the reasons for retraction (Figure 1).

Meta-analyses that included a study retracted for issues with the data, methods or results were significantly more likely to have reported a larger effect sizes (i.e. ESR > 100%), compared to the others (Fisher’s exact test: N=27, P=0.047, Odds ratio=6.91). The magnitude of ESR was significantly different between the two groups, (Wilcoxon rank sum test: W=45, p=0.028). The mean and median ratio values were, respectively, 128.7% and 111.6% for the “data” category and 98.5% and 98.1% for the “other” category.

Despite the low statistical power, estimates calculated on the 12 meta-analyses with inter-transformable data also suggested that articles retracted due to data/methods/results issues (N=5) had inflated the estimated effect size, whereas the others (N=7) had not: the ratio of odds ratio (ROR) was, respectively 1.10 and 1.0 (W=8, p=0.149), the difference in standardized mean difference (DSMD) was, respectively, 0.042 and −0.005, and the difference in correlation coefficient (DCC) was 0.019 and −0.005 (W=7, p=0.106 in both cases).

The number of cases of misconduct in our sample was too small to be meaningfully compared to the rest. Nonetheless, these cases appeared to exhibit similar or larger effects than the others: the average ESR was higher for misconduct-related retractions (mean=138.4) compared to error-related retractions (121.7) and non-data related retractions (98.5), whereas the medians were respectively 104.8, 117.6 and 98.1.

The ratio of CIs was <100 for both groups and not significantly different between them (mean=96.5 and 84.6 respectively, W=71, P=0.373), which suggests that removing any retracted study from a meta-analysis reduced the precision of estimates, regardless of the retraction cause.

### Confounding factors and robustness tests

Meta-analyses with retractions due to data were on average larger than the others, although the difference was not statistically significant (i.e. mean k = 11.4 and 8.3, respectively, W=79 P=0.607). The size of the meta-analysis may represent a confounding factor when comparing the two categories of meta-analyses because larger meta-analyses are likely to be less affected by the removal of any one study (retracted or not) compared to smaller meta-analyses. However, secondary analyses that controlled for the effects of meta-analysis size yielded results of similar or larger magnitude. A multiple linear regression controlling for meta-analysis size suggested that the average ESR was about 30% higher for meta-analyses with retractions due to data, methods or results (b=31.239±15.94, t=1.960, P=0.062). Although the partial effect of retraction type was marginally non-significant, it was much larger than the partial effect of meta-analysis size (b=−0.33±1.10, t=−0.300, P=0.767). An interaction term between the retraction type dummy variable and meta-analysis size was also far from statistically significant (b=−0.87±2.90, −0.299, P=0.768). A partial effect of retraction type was observed if data was weighted by sample size (b=25.57±13.81, t=1.852, P=0.076). If the three most influential points were removed from the multiple regression analysis, to avoid the possible biasing effects of extreme values (see Fig 1), the estimated effect was halved but statistically significant at the 0.05 level (b=13.68±6.01, t=2.28, p=0.034).

As an alternative, more robust and conservative analysis, we run a multiple logistic regression model on the log odds of ESR being larger than 100%. This analysis suggested that, controlling for meta-analysis size, the odds of having ESR >100% were about 50% higher for retractions due to data, methods or results compared to the others (logistic regression: b=0.4±0.18, z=2.182 P=0.039, Odds ratio=1.49). The partial effect of the size of meta-analysis was again far from statistically significant (b=0.01±0.01, z=0.845, P=0.407), nor was there a significant effects for the interaction between meta-analysis size and the reason for retraction dummy variable (respectively, b=−0.03±0.03, z=−0.88, P=0.387). A logistic regression analysis in which observations were weighted by sample size yielded a similar effect size (b=0.41±0.17, t=2.43 P=0.022, Odds ratio=1.5).

If limited to the 12 meta-analyses with inter-transformable data, the multiple linear regression model again confirmed the descriptive data, by suggesting a difference of about 9% in ROR, of 5% in DSMD and 2% in the DCC, although the null hypothesis of no difference was never rejected at the 0.05 level in these analyses, which is not surprising given that sample size are small and thus statistical power is low (df=10, b=0.10±0.07, t=1.466, P=0.177, and b=0.05±0.04, t=1.285, P=0.231, and b=0.02±0.02, t=1.605, P=0.143, respectively).

All results described above where of day excluding four of the eight meta-analyses that shared a retraction. Identical results were obtained if the complementary four meta-analyses were excluded instead.

## DISCUSSION

### Summary of results

Meta-analyses that included a retracted study may or may not have reported an exaggerated effect size, depending on whether the retraction was due to problems with the data, methods or results, as opposed to plagiarism and other non-data-related issues. In the former case, and assuming that the effects reported by the retracted study are entirely incorrect and should be removed, the odds of over-estimating the effect size were 50% larger, and effect sizes were, on average, 30% larger, across effect size metrics. The most conservative estimate in our sample, based on a subsample of N=12 meta-analyses with inter-transformable effect size metrics, suggested that meta-analyses containing a single article with flawed or falsified data reported, on average, 9% larger Odds Ratios, 4% larger Cohen’s d and 2% larger correlation coefficients. Conversely in meta-analyses that included a study that was later retracted for reasons other than problems with data, methods or results, no statistically significant difference of effect sizes was observed.

Two conclusions can be derived from these results, sending a re-assuring and a concerning message, respectively. The re-assuring message is that a sizeable proportion of retractions – namely all retractions that are due to plagiarism or other non data-related breaches of ethics or integrity – are unlikely to have a distorting effect on meta-analyses, and by extension the scientific literature in general. In our sample, retractions of this kind constituted approximately 50% of the data. Estimates in the broader literature vary by discipline and country and are often hampered by incomplete information, but typically suggest that these reasons for retractions account for well over 50% of the total, whereas retractions due to issues with data, methods or results rarely comprise more than 25% of the total (e.g. [17–22].

The troubling message, however, is that retractions due to issues with data, methods or results, regardless of whether those issues were caused by negligence, error or intentional misconduct, may require a downward correction of prior meta-analytical estimates they contributed to. By extension, these results suggest that, regardless of whether a study was included in a meta-analysis, its retraction due problems with data, methods or results might require a downward revision of prior beliefs about the reported phenomenon.

As discussed below, the limitations of this study are unlikely to undermine this key finding. However, as a note of caution, it is worthwhile remarking that in only 1 case out of the 31 cases examined did the retraction lead to an estimate outside the previously calculated 95% confidence interval. Therefore, retractions will not drastically alter the conclusions of meta-analyses if and to the extent that the meta-analysts have taken confidence interval (that is, the precision of their meta-analysis) into account when drawing their conclusions, which is how meta-analytical results ought to be interpreted anyway.

Two limitations may have affected the accuracy of our estimate. The first limitation is a small sample size, which resulted from the choice of sampling recent meta-analyses. Combined with the methodological heterogeneity of meta-analyses (which differed in various characteristics including the statistical metrics used), a small sample size made our estimates rather coarse-grained. Since the nature and magnitude of effects is likely to vary across fields of research, future analyses could repeat our methods in more homogeneous samples of meta-analyses from specific fields.

A second possible limitation in our analysis is the reliance, when classifying the causes of a retraction, on retraction notices. These are known to be an imperfect tool because they may depict the causes of a retraction inaccurately, in particular by under-reporting the occurrence of scientific misconduct [23]. This limitation is likely to weaken our estimate of the difference between retractions due to unintentional errors and those due to misconduct, which was therefore only presented as a secondary and exploratory result. However, this limitation will not affect our main observation of a difference between issues relating to data versus not, because a retraction notice will not misrepresent this aspect (for example, it will not falsely report as plagiarism what was in reality data fabrication). Furthermore, even if this were the case, it would make our present estimate more conservative.

The finding that retractions due to data, methods or results might over-estimate effect sizes, supports the hypothesis that many forms of error, bias and misconduct in science are directed at producing “positive” results, and therefore generally lead to a literature that over-estimates the significance and magnitude of effects. This hypothesis was also supported by a previous meta-meta-analytical study, which found that primary studies whose first authors had articles retracted reported significantly more extreme effect sizes [14]. This hypothesis was also strongly suggested by intuition and experience, leading several authors to assume that bias in research is primarily a bias towards false-positives (e.g. [24]). Therefore, our results confirm previous evidence and intuitive assumptions about a link between (certain kinds of) problematic research and the rate of false positive results in the literature, and offer a preliminary estimate of the magnitude of this effect.

In light of these results, a strong recommendation must be made to reassess and update meta-analyses and possibly other non-quantitative reviews of the literature whenever a retraction is issued, at least when the retraction is due to problems with data, methods or results. This recommendation is of course secondary to that, made repeatedly in the literature, to ensure that retracted studies are not included in meta-analyses to begin with. Clearly, the latter recommendation is still inconsistently followed by meta-analyses who, in our sample, had included studies retracted an average 6 years before their own publication date. In light of our results, neglect to remove retracted studies from meta-analyses can be problematic, at least when retractions are due to problems with data, methods or results.

In conclusion, our findings support concerns that retractions can have a significant impact on the literature, and yet re-affirm the fact, often overlooked, that retractions are not all the same and distinguishing them is important. In line with previous studies on the financial and career impacts of retractions (see Introduction), we found that the epistemic impact of a retraction is significantly linked to the nature of the retraction itself. Therefore, providing clearer and accessible information on the nature of a retraction would not only set fairer incentives for researchers to self-correct their own mistakes [2,3], but it would also help to re-evaluate a scientific literature after errors or misconduct come to light.

## ACKNOWLEDGEMENTS

Julie Wong helped with the retrieval and selection of studies and the collection of most raw data.

## Author contributions

DF: Conceptualization, Data Curation, Formal Analysis, Investigation, Methodology, Project Administration, Resources, Supervision, Validation, Visualization, Writing – Original Draft Preparation. DM: Conceptualization, Validation, Writing – Review & Editing.

## SUPPLEMENTARY INFORMATION

SI1: List of retracted studies, their corresponding retraction, and their corresponding meta-analysis.

SI2: EXCEL file with all primary data and details of all meta-analytical calculations.

SI3: R code of all analyses and figures.

SI4: Raw data used for the analyses.

